# Overexpression of +TIPs EB1, EB3, and DCX in cones of *Danio rerio* results in eye organomegaly and hypertrophy of cone photoreceptors

**DOI:** 10.64898/2026.07.02.736219

**Authors:** Kerstin M Janisch

**Author notes:** ***Abbreviations:*** CMZ, ciliary marginal zone; DCX, doublecortin; dpf, days post-fertilization; EB, end-binding protein; ESCC, esophageal squamous cell carcinoma; IFT, intraflagellar transport; IS, inner segment; MAP, microtubule-associated protein; MMP, matrix metalloproteinase; MT, microtubule; OS, outer segment; RPE, retinal pigment epithelium; RP1, retinitis pigmentosa 1 protein; RP1L1, retinitis pigmentosa 1-like 1 protein; Shh, Sonic hedgehog; +TIP, microtubule plus-end tracking protein.

## Abstract

Photoreceptor outer segments are sensory cilia whose maintenance depends on a balance between basal disc renewal and tip shedding, controlled by intraflagellar transport and axonemal microtubule organization. Microtubule plus-end proteins regulate microtubule dynamics and are strong candidates for roles in this process. In this study, mCherry-tagged EB1, EB3, and DCX were overexpressed in zebrafish (*Danio rerio*) cone photoreceptors under a cone-specific promoter. Eyes were examined at 5 and 10 dpf, and eyecup depth, diameter, and cone photoreceptor area were quantified relative to uninjected controls.

At 5 dpf, all three constructs produced eyes indistinguishable from those of controls. By 10 dpf, all three constructs significantly increased eye cup depth and cone photoreceptor area. EB1 and DCX also significantly increased eye cup diameter. EB1 and, more severely, EB3 also caused retinal holes, mainly in the retinal pigment epithelium and at the outer nuclear/outer plexiform layer, along with misshapen cells near the inner plexiform layer. DXC did not cause retinal holes, but, like EB1 and EB3, produced enlarged, bulbous cone outer segments.

The results show that overexpression of any of the three +TIPs results in a similar eye and photoreceptor overgrowth phenotype, while also producing construct-specific defects: EB1 and EB3 disrupt the broader retinal architecture, whereas DCX produces enlarged eyes. The shared outer segment hypertrophy suggests an imbalance between cargo delivery at the basal end and shedding of the distal tips. The organomegaly may reflect altered progenitor signaling in the ciliary marginal zone.

## INTRODUCTION

Photoreceptors are specialized neuroepithelial cells. Their cell bodies project an axon basally to the synaptic terminal and apically into the inner segments (IS); these IS are connected to the outer segments (OS) by connecting cilia, making the photoreceptor OS specialized primary cilia. The basal body is located at the distal surface of the IS. The OS is stacked with membrane discs containing the vision transduction machinery.

The OS is maintained by a continuous balance between the addition of these discs at the base and the shedding of its distal tip, in accordance with circadian rhythms. The addition of materials at the base depends on the intraflagellar transport (IFT) to deliver structural and signaling proteins along the connecting cilium axoneme and on ciliary ectosomes and endosomal systems. To prevent old discs from accumulating at the tip of the OS, they are released from the tip so that the RPE can engulf and phagocytize them (Young, 1967; Molday and Moritz, 2015; Forrester et al., 2016; Salinas et al., 2017; Carter and Blacque, 2019; Nachury and Mick, 2019; Wensel et al., 2021; Xu, Zhao, and Khang, 2024)

Because OS renewal and maintenance are dependent on a properly organized microtubule (MT) axoneme, the proteins that regulate MT dynamics and IFT cargo delivery are central to photoreceptor structure, function, and survival.

Microtubule plus-end tracking proteins (+TIPs) are a diverse group of proteins that dynamically track and accumulate at the growing plus end of MT. They play a variety of roles, including regulating MT dynamics and stability, anchoring MTs to cellular structures such as the cell cortex, kinetochores, and actin networks, and recruiting additional proteins. End-binding proteins (EB) are well-characterized +TIPs. The EB family consists of evolutionarily conserved proteins: EB1, which is ubiquitously expressed; EB2, which is enriched in the CNS and brain tissue; and EB3, which shows concentrated expression patterns in neural and skeletal tissues. EB1 and EB3 have been shown to be required for cilia assembly and ciliary length regulation. Both proteins localize to the photoreceptor connecting cilium in mammalian cells. (Juwana et al., 1999; Nakagawa et al., 2000; Akmanova and Hoogenraad, 2005; Schrøder et al., 2007; Akhmanova and Steinmetz, 2008; Galjart, 2010; Schrøder et al., 2011; Larsen et al., 2013; Mohan et al., 2013; Hidalgo-de-Quintana et al., 2015; Li et al., 2022; Phillips et al., 2026)

Additionally, EB1 and EB3 have been studied as live-imaging tools for tracking MTs in cultured cells (Stepanova et al., 2003; Komarova et al., 2009), raising the possibility that fluorescently tagged EB constructs could similarly serve as a tool to visualize the dynamics of the photoreceptor axoneme at the distal tip *in vivo*.

The doublecortin family is a family of microtubule-associated proteins (MAPs) and includes two members relevant to photoreceptor biology, RP1 and RP1L1. The family namesake, doublecortin (DCX), itself stabilizes microtubules and modulates kinesin motility by compacting the MT lattice. DCX is essential for neuronal migration and process outgrowth during brain development. DCX is not expressed in the *Danio rerio* retina, whereas RP1 and RP1L1 are present in photoreceptors and required for normal OS disc stacking and axoneme organization. Mutations in RP1 and RP1L1 cause retinitis pigmentosa in humans, and interestingly, overexpression of RP1 also causes retinal degeneration. (Gleeson et al., 1999; Horesh et al., 1999; Liu, Zuo, and Pierce, 2004; Moores et al., 2004; Coquelle et al., 2006; Moores et al., 2006; Jaworski, Hoogenraad, and Akhmanova, 2008; Yamashita et al., 2009; Liu, Zhang, and Pierce, 2010; Liu et al., 2012; Noel and MacDonald, 2020; DeOliveira-Mello et al., 2022)

Because *Danio rerio* lacks endogenous DCX in the retina, it offers a clean slate for modeling the consequences of introducing DCX into photoreceptors. DCX’s use was thought to be a proxy for studying RP1/RP1L1-related dysfunction.

The zebrafish eye offers a particularly useful system for studying questions not only concerning retinal function but also neurogenesis, tissue regeneration, and stem cell niches. Unlike the mammalian eye, the zebrafish eye grows continuously throughout life. New retinal neurons, including photoreceptors, are generated from a ring of stem and progenitor cells at the retinal periphery called the ciliary marginal zone (CMZ). This ongoing growth is tightly regulated by developmental signaling pathways, such as Wnt, Notch, and Sonic hedgehog (Shh). These pathways regulate progenitor proliferation, self-renewal, and differentiation within the CMZ. Several of the proteins examined in this study have documented links to these pathways outside the retina: EB1 binds the tumor suppressor APC, a core component of the Wnt pathway. Dysregulated EB1/EB3 expression has been implicated in promoting proliferation in several cancers, including liver, ESCC, glioblastoma, and gastric carcinoma (EB1) and invasiveness in prostate cancer (EB3). (Johns, 1977; Su et al., 1995; Bart, Siemers, and Nelson, 2002; Kubo, Takeichi, and Nakagawa, 2003; Moshiri, Close, and Reh, 2004; Slep et al., 2005; Wan et al., 2005; Liu et al., 2009; Locker et al., 2006; Lad, Cheshier, and Kalani, 2009; Cerveny, Varga, and Wilson, 2011; Vorday et al., 2012; Fischer, Bosse, and El-Hodiri, 2013; Berges et al., 2014; Richardson et al., 2016; Wan et al., 2016; Dart et al., 2017; Gemoll et al., 2017; Nerli, Rocha-Martins, and Norden, 2020)

This raises the possibility that overexpressing EB1 or EB3 in a continuously growing organ, such as the zebrafish eye, could perturb not only ciliary architecture but also progenitor cell behavior in the CMZ, with possible consequences for eye morphology.

In this study, mCherry-tagged EB1, EB3, and DCX were overexpressed explicitly in zebrafish cone photoreceptors under the cone-specific transducin alpha promoter (Ta-CP) (Kennedy et al., 2007), with the original goal to establish a tool to visualize the distal tip of the cone photoreceptor axoneme. Unexpectedly, this approach revealed that overexpression of any of these three +TIPs produced a phenotype affecting the eye as a whole, characterized by organomegaly and hypertrophy of cone photoreceptors.

This study characterizes the time course, morphology, and severity of these phenotypes for each construct and considers possible mechanisms that could account for these observations.

## MATERIAL AND METHODS

### Animals

All experiments were approved and carried out in accordance with the Institutional Animal Care and Use Committee (IACUC) of the Medical College of Wisconsin. Zebrafish are housed at the zebrafish facility of MCW at 28.5 °C on a 14-hour light and 10-hour dark cycle. Zebrafish were fixed at 5 and 10 dpf after being transferred to a Petri dish containing 0.05% Tricaine in fish water.

### Microinjections

Plasmids of the +TIP constructs were injected into one-to two-cell-stage wild-type embryos using a microinjector (nanoliter 2000 microinjector; World Precision Instruments, Inc). Constructs were diluted in 0.1% Phenol Red (Sigma) to 25 ng/μl (0.3 mM) and injected at 4.6 nl/embryo. Injected and uninjected embryos were kept at 28.5 °C in 0.003% 1-phenyl-2-thiourea (PTU) in fish water to inhibit pigment formation. Fish are screened for expression of mCherry at 5 dpf (Insinna et al., 2008; Insinna et al., 2009a; Insinna et al., 2009b).

### Expression clones +TIP constructs

Invitrogen’s Multisite Gateway® Technology was used to create the constructs for the experiments. EB1, EB3, and DCX PCR products were fused to mCherry, and entry clones with site-specific recombination sites were created. +TIP entry clones and an entry clone containing zebrafish transducin alpha cone promoter (Ta-CP) (Kennedy et al., 2007) were recombined with pcDNA 6.2/V5-pL-DEST vectors to create Ta-CP-EB1mCherry, Ta-CP-EB3mCherry, and Ta-CP-mCherryDCX expression constructs used in the experiments.

### Sample preparation

Embryos were fixed at 5 and 10 dpf in glutaraldehyde/paraformaldehyde (2%/2%) in Sorenson buffer for 2 hours at room temperature. After washing in 0.1 M phosphate buffer, embryos were post-fixed in 1% osmium tetroxide for 1 hour on ice. After dehydration in methanol, embryos are embedded in Embed 812 (Electron Microscopy Sciences) and sectioned at 1 μm using an RMC PowerTome (RMC Boeckeler) (Insinna et al., 2009b).

### Imaging, image processing, and statistics

Ultra-thin sections of embryos were imaged with a TE300 Nikon microscope at 60x. Images were analyzed in Fiji/ImageJ.

For depth and diameter measurements: after setting the scale in Fiji/ImageJ for the objective used to take the pictures, a line was drawn along the distance, and the length was measured with Fiji/ImageJ.

For photoreceptor area measurements: after setting the scale in Fiji/ImageJ for the objective used to take the images, a grid was overlaid on the image (path: ‘Analyze’ tab, ‘Tools’, ‘Grid’). The grid size was set at 0.75 µm^2^ with random offset, and all squares depicting a cone photoreceptor were counted using the point tool of Fiji/ImageJ.

Images of sections close to the optic nerve head were used for all analyses.

Statistical analysis was done in Microsoft Excel. Significance was calculated with Student’s t-test; control sample sizes were n = 5 and n = 7 for zebrafish expressing +TIP clones, respectively.

## RESULTS

### +TIP constructs are expressed in zebrafish eyes

All three mCherry-tagged fusion proteins were expressed exclusively in the zebrafish eyes. At 5 dpf, the eyes exhibit a robust fluorescent signal well above background autofluorescence (Figure 1, A-C). The eyes look normal and healthy in DIC pictures, and overlays indicate that the mCherry fluorescence is only contained in the eyes (Figure 1, A’-C’).

**Figure 1:**
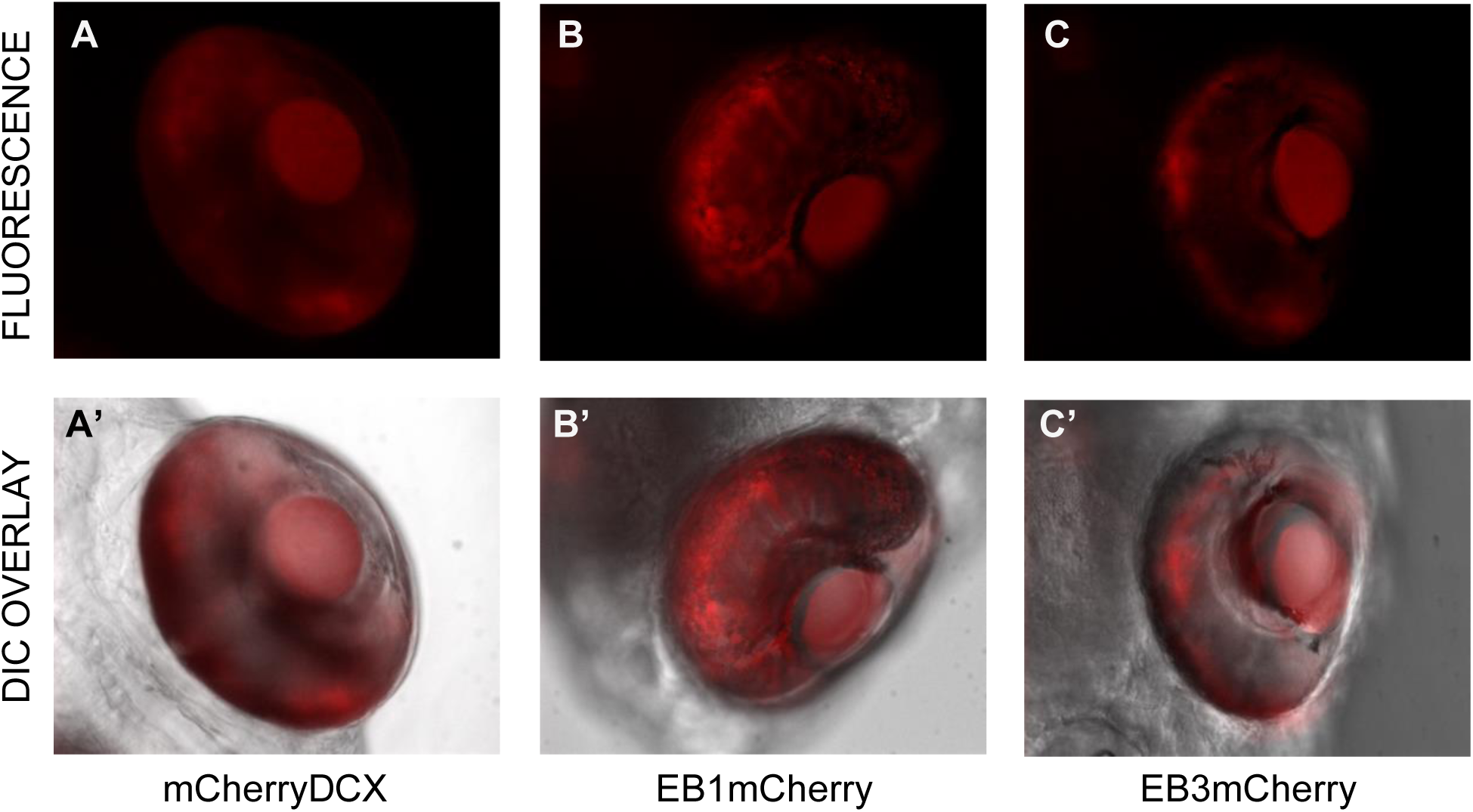
Representative fluorescence and DIC images of 5 dpf zebrafish eyes expressing mCherry-tagged fusion proteins. A, fluorescence of zebrafish eye expressing mCherryDCX and A’, fluorescence and DIC overlay of zebrafish eye expressing mCherryDCX; B, fluorescence of zebrafish eye expressing EB1mCherry and B’, fluorescence and DIC overlay of zebrafish eye expressing EB1mCherry; C, fluorescence of zebrafish eye expressing EB3mCherry and C’, fluorescence and DIC overlay of zebrafish eye expressing EB3mCherry

### Eye morphology is normal at 5 dpf

To assess any possible influence of the overexpression of the three fusion proteins in the zebrafish cones, osmium-postfixed eyes were cross-sectioned and imaged at 20x and 60x (Figure 2). At 20x, gross eye morphology of eyes expressing either EB1mCherry, EB3mCherry, or mCherryDCX (Figures 2B, 2C, and 2D, respectively) presents as normal when compared to control eyes (Figure 2A). Tears and holes in the lens are artifacts from fixation and cutting. All eyes show correct stratification of the retinal layers, as confirmed by images at 60x magnification. All eyes have properly formed cone photoreceptors (Figure 2, A’-D’).

**Figure 2:**
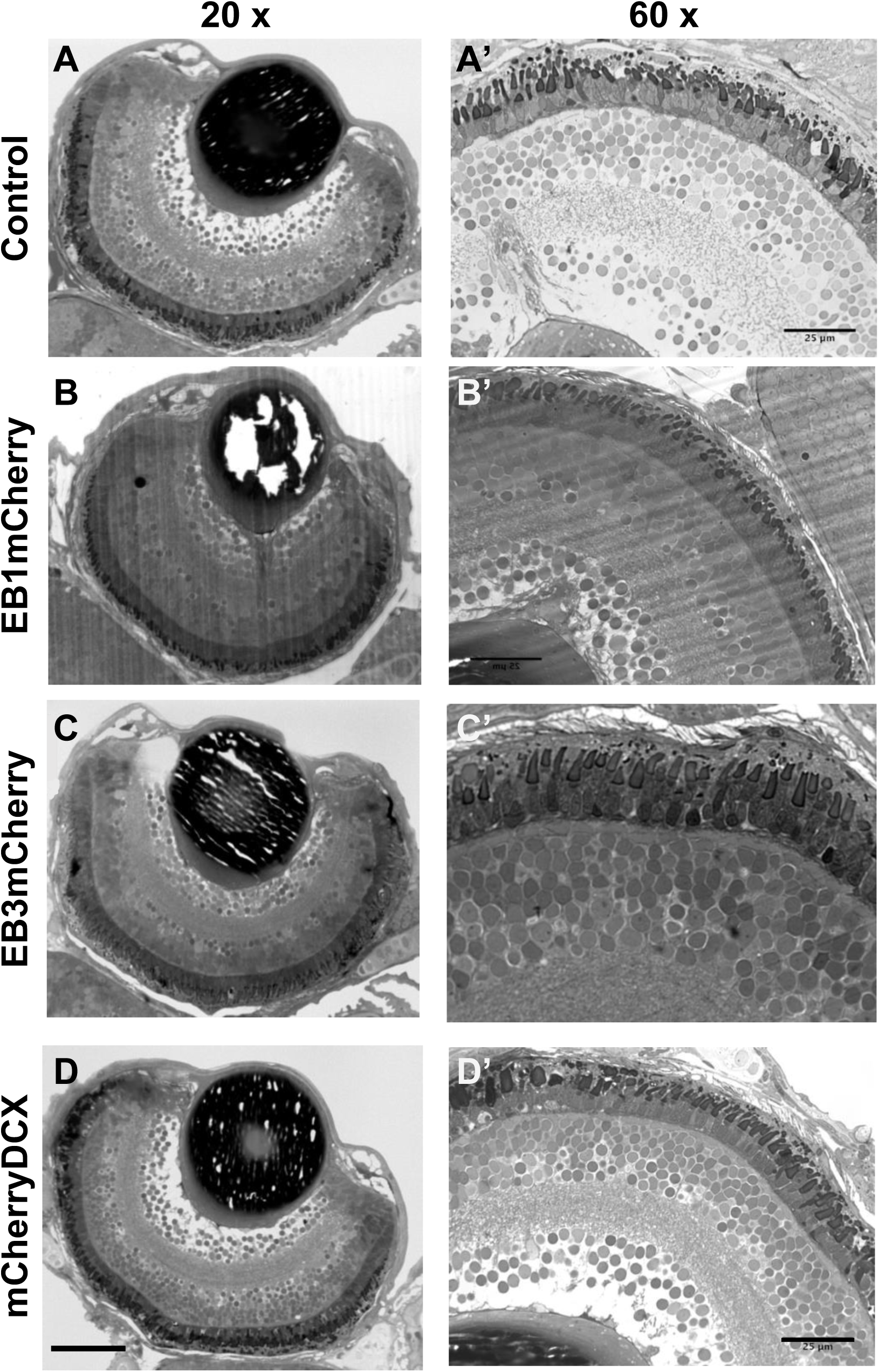
Cross-section of eye morphology of zebrafish at 5 dpf at 20x and 60x magnification. A, morphology of a representative control eye of uninjected control at 20x; A’, 60x magnification of retina of control eye; B, morphology of a representative eye of a zebrafish injected with EB1mCherry; B’ 60x magnification of retina of EB1mCherry injected zebrafish; C, morphology of a representative eye of a zebrafish injected with mCherryDCX; C’, 60x magnification of retina of EB3mCherry zebrafish; D, morphology of a representative eye of a zebrafish injected with EB3mCherry; D’, 60x magnification of retina of mCherryDCX zebrafish. The scale bar represents 50 μm for images A – D and 25 μm for images A’ – D’

### Fusion proteins disrupt eye morphology at 10 dpf

At 10 dpf, the eyes show clear signs of disrupted eye morphology. Eyes injected with either EB1mCherry or EB3mCherry (Figure 3, B and C, respectively) appear to have a deeper eye cup depth when compared with the control (Figure 3, A). The eyes expressing either EB-fusion proteins show holes of various sizes between the outer nuclear and outer plexiform layer (Figures 3B and C, respectively). Whereas eyes expressing mCherryDCX are larger in diameter but do not seem to have developed holes (Figure 3D).

**Figure 3:**
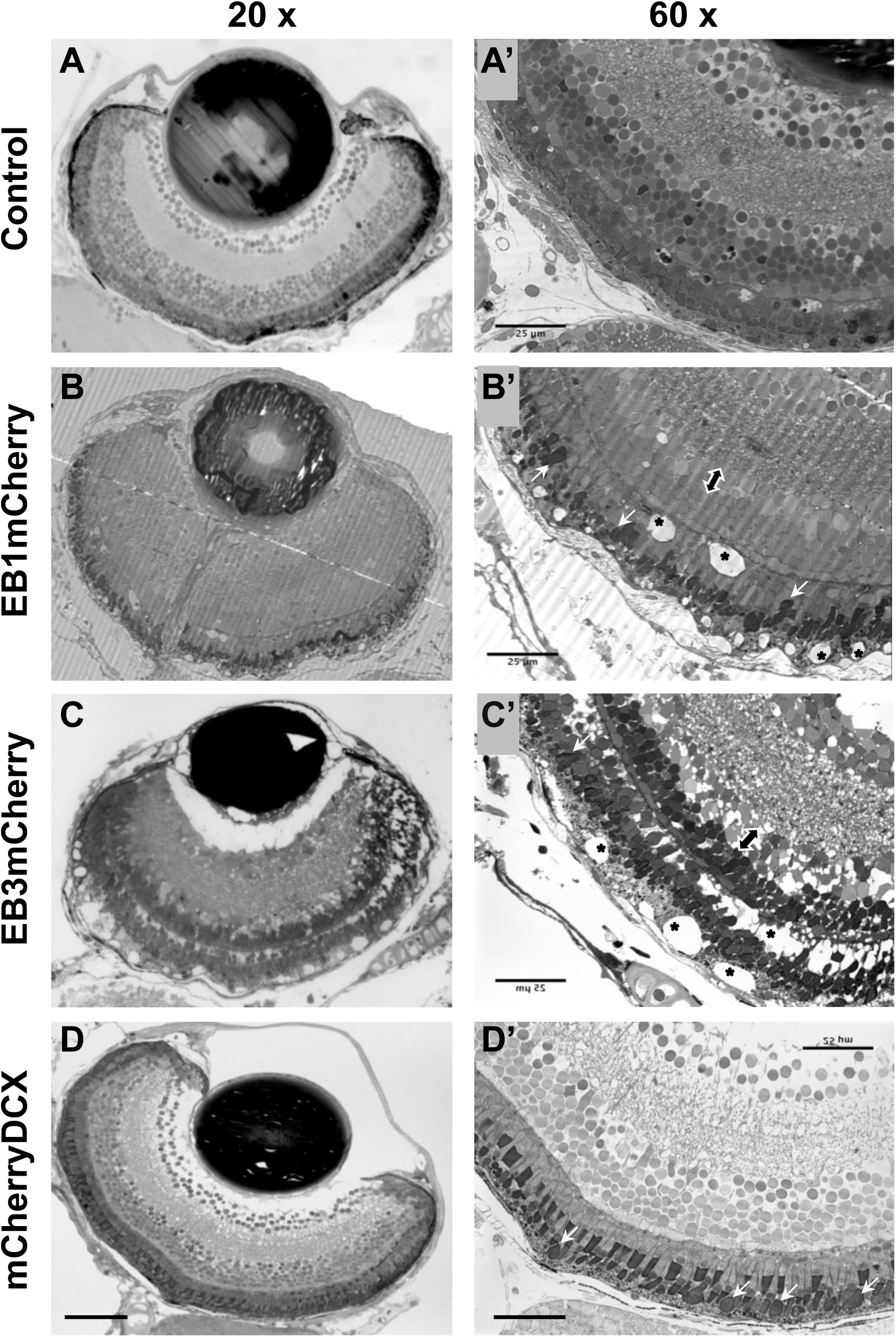
Cross-section of eye morphology of zebrafish at 10 dpf at 20x and 60x magnification. A, morphology of a representative control eye of an uninjected control at 20x; A’, 60x magnification of retina of control eye; B, morphology of a representative eye of a zebrafish injected with EB1mCherry; B’ 60x magnification of retina of EB1mCherry injected zebrafish; C, morphology of a representative eye of a zebrafish injected with mCherryDCX; C’, 60x magnification of retina of EB3mCherry zebrafish; D, morphology of a representative eye of a zebrafish injected with EB3mCherry; D’, 60x magnification of retina of mCherryDCX zebrafish. The scale bar in D represents 50 μm for images A – D and in D’ 25 μm for images A’ – D’ * mark holes in retinal layers; white arrows mark misshapen cone photoreceptors; double-headed arrows mark nuclear rows with misshapen cells

At 60x magnification, it becomes clear that the overexpression of either of the two EB-fusion proteins not only resulted in holes within the retinal layers (asterixis in Figure 3B’ and C’) but also in misshapen cells in the lower layers of the inner nuclear layer bordering on the inner plexiform layer (double-headed arrows in Figure 3B’ and C’). The holes primarily occur within the RPE, but gaps can also be found between the outer nuclear layer and the outer plexiform layer. Overexpression of EB3mCherry results in a more severe phenotype than EB1mCherry overexpression. Some cone photoreceptors appear to be increased in size, misshapen, and bulbous (white arrows in Figure 3B’ and C’). mCherryDCX-expressing eyes exhibit nice retinal layering without holes, but several cone photoreceptors appear to be larger, misshapen, and bulbous (white arrows in Figure 3D’).

### Quantification of eye cup depth and diameter at 5 and 10 dpf

Quantification of eye cup depth and diameter revealed no differences for any of the used constructs at 5 dpf (Figure 4, A and D). However, at 10 dpf, there is an apparent increase in eye cup depth for all constructs used. This increase is statistically significant for all three constructs (Figure 4B). Regarding eye cup diameter, overexpression of mCherryDCX results in a highly statistically significant increase at 10 days post-fertilization (dpf). For the two EB constructs, only EB1mCherry overexpression induces a significant increase in eye cup diameter, whereas EB3mCherry is trending toward a larger eye cup diameter but does not reach statistical significance (Figure 4E). Figures 4C and F show representative images used for quantification, with arrows indicating the measurement line.

**Figure 4:**
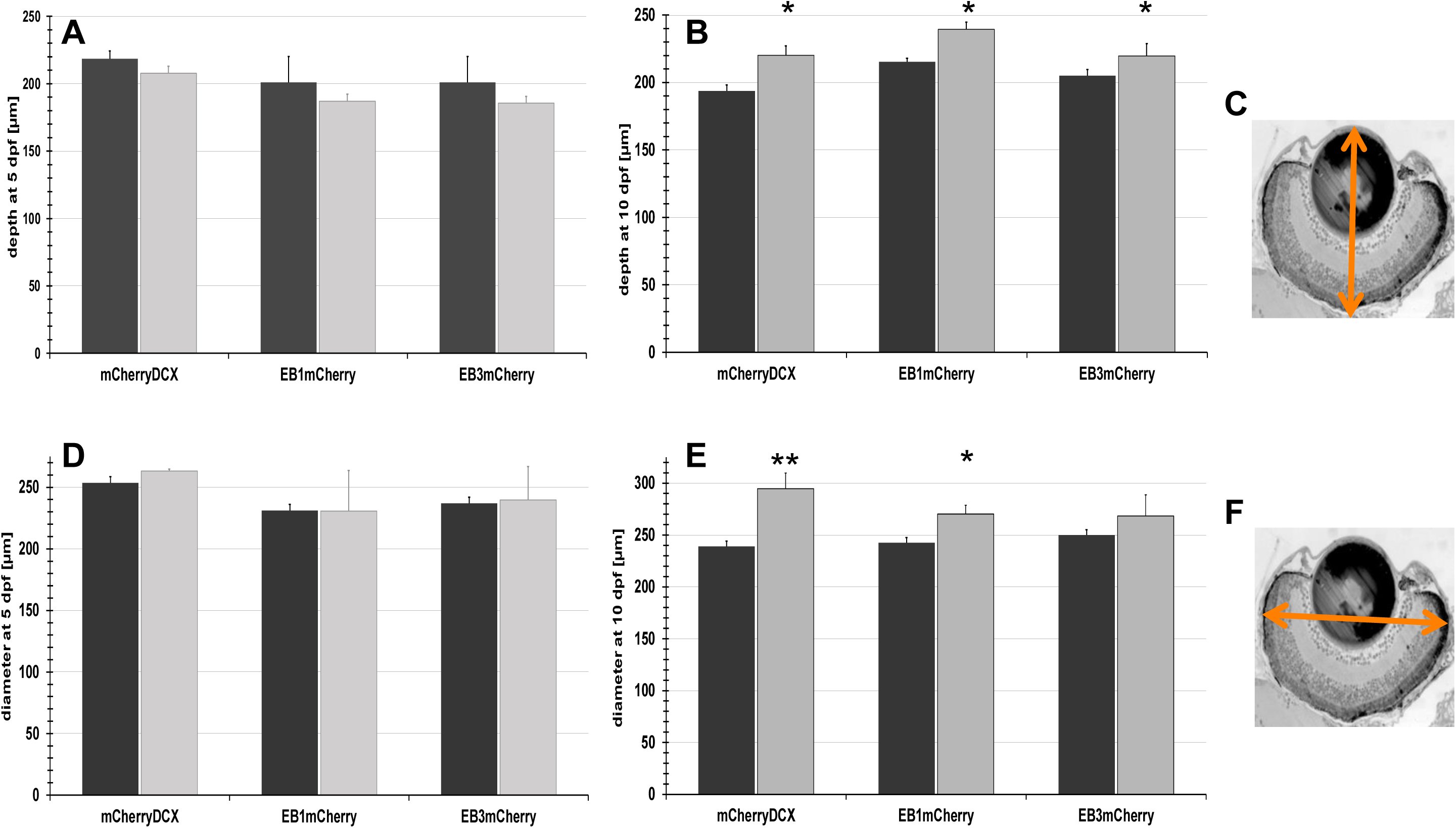
Measurements of eyecup depth and diameter at 5 dpf and 10 dpf. A, measurements of eyecup depths of transfected zebrafish with mCherryDCX, EB1mCherry, and EB3mCherry and their respective controls at 5 dpf; B, measurements of eyecup depths of transfected zebrafish with mCherryDCX, EB1mCherry, and EB3mCherry and their respective controls at 10 dpf; C, picture of a representative eye section to show the path of measurement (orange arrow); D, measurements of eyecup diameter of transfected zebrafish with mCherryDCX, EB1mCherry, and EB3mCherry and their respective controls at 5 dpf; E, measurements of eyecup depths of transfected zebrafish with mCherryDCX, EB1mCherry, and EB3mCherry and their respective controls at 10 dpf; F, picture of a representative eye section to show the path of measurement (orange arrow); error bars are stdev; student t-test was used to calculate significance; * p ≤ 0.05, ** p ≤ 0.01; n = 5 for all controls, n = 7 for all transfects.

### Quantification of cone photoreceptor area at 5 and 10 dpf

Considering all three constructs were created under the transducin alpha cone promoter, the cone photoreceptors ought to be the cells containing most, if not all, of the expressed protein. It is sensible to presume that they are the most affected. Quantification at 5 dpf reveals no statistically significant increase in cone photoreceptor area but indicates a trend toward an increase (Figure 5A). At 10 dpf, all three constructs produce highly statistically significant increases in cone photoreceptor area (Figure 5B). The image in Figure 5C shows a representative image used for quantification, along with an exemplary grid (not to scale).

**Figure 5:**
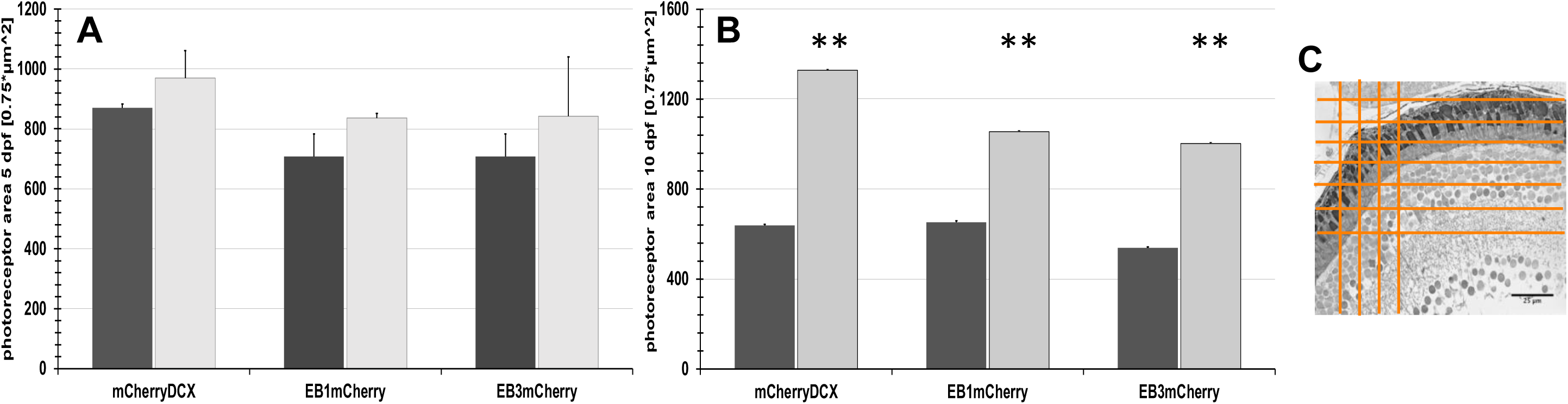
Measurements of cone photoreceptor area of transfected zebrafish at 5 dpf and 10 dpf. A, measurements of photoreceptor area of transfected zebrafish with mCherryDCX, EB1mCherry, and EB3mCherry and their respective controls at 5 dpf; B, measurements of photoreceptor area of transfected zebrafish with mCherryDCX, EB1mCherry, and EB3mCherry and their respective controls at 10 dpf; C, representative picture of a 60x section of a retina with an exemplary grid; error bars are stdev; student t-test was used to calculate significance; ** p ≤ 0.01; n = 5 for all controls, n = 7 for all transfects

## DISCUSSION

In this work, mCherry-tagged EB1 and EB3, as well as DCX, were overexpressed in the cone photoreceptors of *Danio rerio* under the Ta-CP promoter. The +TIPs EB1 and EB3 have been used as tools to track MT dynamics *in vitro* (Stepanova et al., 2003; Ma et al., 2004; Akhmanova and Steinmetz, 2008; Komarova et al., 2009; Galjart, 2010; Nehlig et al., 2017). The DCX construct was chosen as a proxy of RP1 and RP1L1 (retinitis pigmentosa 1 and retinitis pigmentosa 1-like 1 protein, respectively), both of which belong to the doublecortin family. (Liu et al., 2003; Liu, Zuo, and Pierce, 2004; Reiner et al., 2006; Noel and MacDonald, 2020)

The author set out to establish a system to track and visualize the distal tip of the cone photoreceptor axoneme for further studies. Interestingly, the overexpression of these +TIPs resulted in phenotypes characterized by eye organomegaly and hypertrophy of the cone photoreceptors.

### Eye Phenotype by Overexpression of EB1 and EB3

Overexpression of EB1 and EB3 resulted in a change in the eye morphology and in a quantifiable increase of eye cup depth and photoreceptor area at 10 dpf (Figures 3, 4B and E, and 5B). At an earlier time point, 5 dpf, eye morphology, eye cup depth, and diameter were normal (Figure 2, 4A and D), and photoreceptor area showed a non-significant trend towards an increase (Figure 5A) when compared to controls. Interestingly, the EB3 overexpression phenotype was more severe than that of EB1 overexpression (Figure 2). Both EB constructs result in retinal holes, predominantly within the RPE layer and between the outer nuclear and outer plexiform layers (Figure 3). The retinal tissue expressing the EB3-fusion protein appears to have a looser cell-cell formation and increased interstitial or extracellular space (Figure 3C’). Another phenotype of the overexpression of the two EB-fusion proteins is the appearance of misshapen cells within the lower layers of the inner nuclear layer bordering on the inner plexiform layer (Figure 3, B’ and C’, double-headed arrow). A closer look at the cone photoreceptors exhibits misshapen and bulbous cone outer segments (white arrows in Figure 3, B’ and C’). The quantification of the photoreceptor area shows that both EB-fusion proteins result in a highly statistically significant increase in the photoreceptor area at 10 dpf (Figure 5B).

EB1 is expressed ubiquitously in tissues, whereas EB3 exhibits a more specific expression pattern, with a concentration in nervous and skeletal tissues. Both EBs are vital proteins for cilia formation and length (Schrøder et al., 2007; Tirnauer and Bierer, 2000; Nakagawa et al., 2000; Schrøder et al., 2011) and have been shown to bind to APC (adenomatous polyposis coli tumor suppressor), an important component of the Wnt pathway (Su et al., 1995; Barth, Siemers, and Nelson, 2002; Slep et al., 2005; Morrison, 2009).

With the usage of the Ta-CP promoter to express the EB-fusion proteins, the expression of both constructs is restricted to the cone photoreceptors (Kennedy et al., 2007). As the zebrafish aged, the author expected to observe morphological changes in the photoreceptor, primarily an increase in outer segment length, given that EB3 overexpression increases cilia length in RPE cells (Schrøder et al., 2011). The ocular organomegaly resulting from fusion protein overexpression is noteworthy. The increase in cone photoreceptor area can be attributed to the accumulation of the EB-fusion proteins in the cilia. Both EB1 and EB3 regulate MT dynamics by binding to the growing plus end of the MT. Schrøder et al. (2011) reported that EB3 overexpression in RPE cells led to elongated cilia, suggesting EB3 plays a role in axoneme elongation. EB1 binds with the tail of the motor protein Kif17 and enhances its ability to bind to MT. This EB1/Kif17-tail complex can modulate Kif17’s regulatory function on MT stability (Jaulin and Kreitzer, 2010; Acharya, Espenel, and Kreitzer, 2013). Kif17 is an essential motor protein found in zebrafish photoreceptor outer segments (Insinna et al., 2008; Lewis et al., 2017; Lewis et al., 2018).

The zebrafish retina’s specialty of continuously producing all retinal cell types throughout the fish’s life is due to the ciliary marginal zone (CMZ). This circumferential ring of retinal stem and progenitor cells is located at the retina’s periphery. The CMZ is responsible for the continuous growth and expansion of the zebrafish eye throughout its lifespan (Johns, 1977; Moshiri, Close, and Reh, 2004; Cerveny, Varga, and Wilson, 2011; Fisher, Bosse, and El-Hodiri, 2013; Wan et al., 2016). Overexpression of EB-fusion proteins in newly forming cone photoreceptors within the CMZ could lead to the translocation of APC from the multi-protein complex comprising axin, GSK3, CKI, and β-catenin. The binding of EB to APC would lead to the complex’s dissociation and the accumulation of β-catenin in the cytoplasm. β-Catenin then moves into the nucleus and activates TCF/LEF transcription factors, which promote transcription of cell proliferation and growth genes. The transcription of Wnt-dependent genes could explain the increase in the eye organomegaly. The dissociation of β-catenin from the multiprotein complex and its nuclear translocation could reduce β-catenin at cell junctions, leading to gaps within the tissue (retinal holes). (Kubo, Takeichi, and Nakagawa, 2003; Lad, Cheshier, and Kalani, 2009; Liu et al., 2009; Morrison, 2009; Borday et al., 2012; van der Wal and van Amerongen, 2020)

Another possibility for the tissue phenotype could be the transcription of metalloproteinases (MMP), which are β-catenin-regulated genes (Wu B., Crampton S.P., and Hughes C.C.W., 2007; Ge et al., 2009; Ingraham et al., 2011). Depending on the MMP, its expression could perpetuate the stem cell population in the CMZ, leading to increased cell production and, consequently, increased eye size (Chang and Werb, 2001; Visse and Nagase, 2003; Kessenbrock, Wang, and Werb, 2015). MMPs could also be degrading extracellular matrix components, leading to edema. It has been shown that MMPs contribute to diabetic retinopathy and the breakdown of the integrity of the blood-retinal barrier (Kowluru and Mishra, 2017; Cui, Hu, and Khalil, 2017; Opdenakker and Abu El-Asrar, 2019; Haydinger et al., 2023). MMPs degrade VE-cadherin in diabetes, and MMP inhibition increases tight junction function (Alexander and Elrod, 2002; Akahane et al., 2004; Navaratna et al., 2007).

When considering the possibility of MMP expression in the CMZ, upregulation could affect the Notch pathway. The Notch pathway is highly conserved and plays a role in organ development by promoting self-renewal and dedifferentiation of stem and progenitor cells (Livesey and Cepko, 2001; Nerli, Rocha-Martins, and Norden, 2020; Zhou et al., 2022; Gozlan and Sprinzak, 2023; Guerin et al., 2025). MMPs can act upstream of the Notch pathway by activating or cleaving receptors and ligands, which could explain the organomegaly of the eyes (Sawey and Crawford, 2008; Zhou et al., 2022).

Another aspect to consider is the Sonic Hedgehog (Shh) pathway, a key signaling regulator in the CMZ. Shh controls cell-cycle exit and regulates cell proliferation, thereby promoting progenitor differentiation (Locker et al., 2006; Borday et al., 2012). To date, there has been no direct link between EB proteins and the Shh pathway; however, the overexpression of EB1 and EB3 could potentially dysregulate this pathway, contributing to the observed phenotype.

### Eye Phenotype by Overexpression of DCX

Overexpression of DCX did not alter the eye morphology, depth, and diameter (Figure 2D and D’) at 5 dpf. At 10 dpf, the eyes still exhibit normal morphology, but 60x pictures reveal bulbous and misshapen cone outer segments (Figure 3D’, white arrows). It also led to a significant increase in eye cup depth and diameter at 10 dpf (Figures 4B and 4E). Quantification of photoreceptor area reveals a trend toward increased area at 5 dpf (Figure 5A), but this difference is not statistically significant. The photoreceptor area is highly statistically increased at 10 dpf (Figure 5B).

DCX is an MT-associated protein (MAP) and the namesake of the DCX family, of which RP1 and RP1L1 are also a part. It is crucial for neuronal development, particularly in processes such as neuronal migration, positioning, and process outgrowth in the brain and retina (Gleeson et al, 1999; Lee et al., 2003; Coquelle et al., 2006; Reiner et al., 2006; Jaworski, Hoogenraad, and Akhmanova, 2008; Yamashita et al, 2009; DeOliveira-Mello et al., 2022). DCX stabilizes straight MT and, by compacting the MT lattice, influences kinesin motility (Horesh et al, 1999; Moores et al., 2004; Moores et al., 2006; Moslehi, Ng, and Bogoyevitch, 2017). It also binds to kinesin-3 motors, modulating their microtubule binding affinity (Moores et al., 2006).

DCX is not expressed in photoreceptors in the mature retina but is present in neuronal cells in the inner nuclear and outer plexiform layer in adult rats (Wakabayashi et al., 2008). It is found in photoreceptor cells in the sea lamprey, in both larval and adult retinas (Fernandez-Lopez et al., 2016). DCX is absent in *Danio rerio* retinas (DeOliveira-Mello et al., 2022) and is used here as a stand-in for RP1 and RP1L1. While RP1 mutations cause retinitis pigmentosa, RP1 overexpression in photoreceptors also leads to retinal degeneration (Liu, Zao, and Pierce, 2004; Liu, Zhang, and Pierce, 2010; Liu et al., 2012; Noel and MacDonald, 2020). The overexpression of DCX in the photoreceptors does not show signs of retinal degeneration, but rather normal, albeit larger, eyes (Figures 2D and D’, 3D and D’, 4). DCX has not been linked to binding any proteins in the Wnt pathway. The increase in eye cup diameter and depth could result from dysregulation of a different developmental pathway in the CMZ, such as the Shh, Hippo, or Notch pathways.

### Photoreceptor phenotype of overexpression of EB1/EB3 and DCX

Interestingly, in all three fusion protein-expressing eyes, deformed, bulbous photoreceptors are found throughout (white arrows in Figure 3, B’, C’, and D’). The deformed, bulbous photoreceptors point to a defect in the organization of outer segment shape and size.

Outer segments maintain their length by balancing disc renewal at their base and disc shedding at the distal end, as well as through IFT-dependent cargo delivery of structurally necessary proteins and vesicular trafficking (Perkins and Fadool, 2010; Molday and Moritz, 2015; Kocaoglu et al., 2016; Salinas et al., 2017; Wensel et al., 2021; Xu, Zhao, and Kang, 2024).

EB1 and EB3 are localized in the connecting cilium of the photoreceptor in mouse and human cells (Schrøder et al., 2011; Hidalgo-de-Quintana et al., 2015). Both play a role in cilia formation. Schrøder et al. (2011) showed that they are important for anchoring minus-end MT to the basal body/centrosome and for facilitating vesicular trafficking.

The bulbous deformation of the outer segment could therefore indicate an imbalance between tip shedding and renewal at the base. Overexpression of both EBs in the outer segment might increase the influx of IFT-dependent cargo, and, at a normal rate of tip shedding, material would accumulate within the outer segment, leading to its deformation.

DCX is found in the outer segments of photoreceptors of lampreys at different developmental stages (Fernandez-Lopez et al., 2016). As mentioned earlier, DCX is not expressed in *Danio rerio*, but RP1 and RP1L1, both from the DCX family, are expressed in the eye and are involved in photoreceptor development (Reiner et al., 2006; Liu, Zhang, and Pierce, 2010; Noel and MacDonald, 2020; DeOliveira-Mello et al., 2022). In *Drosophila*, a DCX-related protein, DCX-EMAP, has been identified as essential for MT organization in ciliary dilations within mechanoreceptors (Bechstedt et al., 2010). The authors hypothesize that DCX-EMAP forms a protein network that links other proteins to the MT of sensory cilia. The overexpressed DCX in this study could also fulfill this role by binding proteins to the axoneme, resulting in material accumulation and bulging of the outer segment.

### Hypothetical considerations and future directions

#### EB1/EB3 eye morphology phenotype

As discussed above for the EB1/EB3 phenotype, the interactions between EB proteins and APC, as well as β-catenin translocation, should be investigated throughout the retina, including the CMZ. The author predicts that differences in the severity of the retinal hole phenotype might be due to EB1 binding to Kif17, which is present in photoreceptors. The binding of EB1 to Kif17 would reduce the available pool of EB1 for APC binding. This could lead to a less severe phenotype than observed with EB3 overexpression, since no interaction between EB3 and Kif17 has been reported. Using Kif17 morpholinos to knock down the kinesin should result in a phenotype similar to that of EB3 overexpression. It would be valid to investigate the Wnt pathway in the CMZ and assess the transcription of Wnt-dependent gene products. Given the phenotype, MMP expression, and the Notch pathway warrant investigation.

Although EB proteins are not considered part of the Shh pathway, given its major role in the CMZ, it is worthwhile to investigate a possible dysregulation of the Shh pathway.

#### DCX eye morphology phenotype

Interestingly, overexpression of DCX results in normal eye morphology, albeit with larger eye cup depth and diameter, as evidenced by quantification. It would be interesting to quantify each retinal layer by thickness and the number of nuclear rows to determine whether any layers contribute to the increased eye size. As mentioned before, DCX is not expressed in the retina of *Danio rerio* but is found in horizontal cells and retinal ganglion cells in mammals (Lee et al., 2003; Wakabayashi et al., 2008; Fernandez-Lopez et al., 2016; DeOliveira-Mello et al., 2022). Therefore, quantifying these cell types in DCX-overexpressing eyes might be warranted.

Another aspect to consider is DCX’s role in glioblastoma. Generally considered a cytoplasmic protein, DCX translocates to the nucleus in glioma cells, contributing to tumor invasiveness and the proliferation and self-renewal of glioblastoma stem-like cells (Ayanlaja et al., 2017; Ayanlaja et al., 2020). This potential for proliferation and self-renewal could also explain the increased eye size and merits further investigation of DCX localization within the retina and the CMZ.

#### Photoreceptor phenotype

Overexpression of EB1/EB3 and DCX leads to the formation of bulbous, deformed cone photoreceptor outer segments. The bulbous deformation appears to indicate an accumulation of material within the photoreceptor’s outer segment. This can be due to an imbalance in the equilibrium between disc shedding at the distal tip and material addition at the base.

A closer look at the outer segment morphology could provide more insight. For the EB1/EB3 constructs, investigating IFT-dependent cargo transport and vesicle formation at the base might also shed more light on outer segment deformations. At the same time, the shed and RPE-engulfed disc packages from the tips could be quantified for each construct to better understand the possible influence of the overexpressed proteins on shedding.

If the DCX construct forms protein networks that link other proteins to the axoneme, as suggested by Bechstedt et al. (2010) for DCX-EMAP, a closer look at outer segment morphology using electron micrographs, along with the use of altered DCX constructs that cannot bind to MTs, can help shed light on this result.

### Conclusions

This work shows that the overexpression of +TIPs in *Danio rerio* cone photoreceptors causes eye organomegaly and a photoreceptor hypertrophy phenotype. The effects become pronounced and statistically significant by 10 dpf for all three fusion proteins. This suggests a developmental or cumulative effect.

EB1/EB3 and DCX overexpression produce overlapping and distinct phenotypes. Both EB proteins cause organomegaly, photoreceptor outer segment deformation, and structural retinal disorganization (EB3 more severe than EB1), whereas DCX causes organomegaly and photoreceptor hypertrophy but no retinal holes. This suggests a perturbation of corresponding but mechanistically distinct pathways.

A common phenotype is the presence of deformed outer segments of photoreceptors, which may reflect disrupted outer segment renewal dynamics. All three +TIP fusion proteins cause bulbous, misshapen outer segments, implying an imbalance between IFT-dependent material delivery at the base and disc shedding at the tip.

## ACKNOWLEDGMENTS

The author would like to thank Dr. Besharse for providing lab space, materials, and instruments at the Medical College of Wisconsin, as well as for mentorship on this work. The funding for this work came from an NIH grant and an NIH core grant for vision research.

The author would also like to thank Dr. Link for access to the Medical College of Wisconsin fish facility.

A big thank you goes to Dr. Willardsen and Pat Cliff for their help with zebrafish husbandry and microinjections.

